# Tactile spatial discrimination on the torso using vibrotactile and force stimulation

**DOI:** 10.1101/2021.03.26.437195

**Authors:** Atena Fadaei J., Matteo Franza, Oliver Alan Kannape, Masayuki Hara, Olaf Blanke

## Abstract

There is a steadily growing number of mobile communication systems that provide spatially encoded tactile information to the humans’ torso. However, the increased use of such hands-off displays is currently not matched with or supported by systematic perceptual characterization of tactile spatial discrimination on the torso. Furthermore, there are currently no data testing spatial discrimination for dynamic force stimuli applied to the torso. In the present study, we measured tactile point localization (PL) and tactile direction discrimination (DD) on the thoracic spine using two unisex torso-worn tactile vests realized with arrays of 3×3 vibrotactile or force feedback actuators. We aimed to, firstly, evaluate and compare the spatial discrimination of vibrotactile and force stimulations on the thoracic spine and, secondly, to investigate the relationship between the PL and DD results across stimulations. Thirty-four healthy participants performed both tasks with both vests. Tactile accuracies for vibrotactile and force stimulations were 60.7% and 54.6% for the PL task; 71.0% and 67.7% for the DD task, respectively. Performance correlated positively with both stimulations, although accuracies were higher for the vibrotactile than for the force stimulation across tasks, arguably due to specific properties of vibrotactile stimulations. We observed comparable directional anisotropies in the PL results for both stimulations; however, anisotropies in the DD task were only observed with vibrotactile stimulations. We discuss our findings with respect to tactile perception research as well as their implications for the design of high-resolution torso-mounted tactile displays for spatial cueing.

## Introduction

As the human torso provides an extensive skin area to convey tactile information, torso-worn haptic displays deploying tactile spatial cues have gained increasing attention in recent years (Rupert 2000; Lemmens et al. 2009; Arafsha et al. 2015; Lentini et al. 2016; Wacker et al. 2016; Buimer et al. 2018; Garcia-Valle et al. 2018). Moreover, while providing tactile information on the torso, a person’s active body parts, such as hands and fingers, remain fully available for daily living activities. Accordingly, the torso has been considered as one of the most available and practical candidate sites for wearable and mobile tactile communication systems (Cholewiak et al. 2004; Kristjánsson et al. 2016) and may also be particularly suited for applications in cognitive and clinical neurosciences (Rognini and Blanke 2016). However, to effectively convey spatially encoded tactile information and make use of this information, more data about the tactile spatial resolution of the torso are required. Surprisingly, we lack such information: even though psychophysical research on tactile spatial acuity was launched in the nineteenth century by Ernst Heinrich Weber (1834), since then, only a handful of studies has studied tactile perception on the torso. In the present study, we described new systems and focused on assessing tactile spatial discrimination of the human torso.

Tactile point localization (PL), evaluates a person’s ability to localize the point of tactile stimulation in an array of stimulators mounted on the torso. Previous studies that used a linear one-dimensional array of vibrators around the waist reported PL accuracies within the range of 74% (12-tactor with 72 mm spacing) to 98% (8-tactor with 107 mm spacing). It was noted that accuracy tended to increase by increasing inter-stimulator spacing and at locations closer to the body midline (i.e., navel, spine) (Cholewiak et al. 2004). The higher localization ability in proximity to specific anatomical reference points, such as the body midline (e.g., navel and spine) and joints (e.g., wrist), was first described by Weber (1834) and was recently confirmed for the localization of both vibrotactile and static pressure stimuli presented on the upper limbs (e.g., wrist) (Cholewiak and Collins 2003; Oakley et al. 2005). However, studies employing two-dimensional arrays of vibrators (e.g., 4×4 array) have not found higher PL accuracy for midline regions and rather observed that PL accuracy changes depending on the location of the target within the array (Lindeman and Yanagida 2003; Cholewiak and McGrath 2005; Jones and Ray 2008). For instance, accuracy was found to vary strongly (from 40 to 82%) depending on the position of the vibrator in the 4 x 4 array (Lindeman and Yanagida 2003; Cholewiak and McGrath 2005; Jones and Ray 2008).

Several other studies have investigated the spatial acuity for tactile cues applied to the torso and measured the capability of discriminating two nearby tactile stimuli presented on the skin surface. Although classical studies used the two-point discrimination test (Weber 1834), more recent studies have questioned the validity of this measure as it is vulnerable to several possible confounds (e.g., two-point discrimination may be based on intensity rather than spatial cues). Alternatively, they suggested a task in which two successive stimuli are applied at nearby locations, and participants’ have to judge whether the two stimuli were delivered in the same location or not (Johnson and Phillips 1981; Johnson 1994; Stevens and Patterson 1995; Tong et al. 2013). For instance, Eskildsen et al. (1969) presented successive vibrotactile stimuli via a horizontal array of five vibrators on the back and reported a discrimination threshold at 10 mm on the back (at the level of the scapula). Van Erp (2005a) measured tactile direction discrimination (DD) using two successive tactile stimuli on the torso, defined as the ability to discriminate whether a second tactile stimulus was to the left or to the right of a first tactile stimulus. They used a linear array of vibrators (11 in horizontal and 14 in the vertical direction). Using this method, they determined the tactile spatial acuity threshold at 20-30 mm on the torso, with better DD accuracy (approximately 10 mm) only for horizontal array locations near to the body midline (i.e., navel and spine). They also highlighted the role of spatiotemporal factors by observing that the accuracy increased as the burst duration and/or inter-stimulus interval increased. Finally, Jóhannesson et al. (2017) explored the impact of inter-stimulator distancing on tactile DD accuracy (so-called relative spatial acuity; three-alternative force choice task [AFC]), using arrays of 3×3 vibrators on the lower thoracic region of the back. They reported that accuracy increased from 64% to 91% as an inter-stimulator spacing increase from 13 to 30 mm. Taken together, while PL and DD measure two different aspects of tactile spatial discrimination, previous studies often directly compared the results of these two tasks. To the best of our knowledge, there is no study investigating the degree of agreement or disagreement between the results of tactile localization and direction discrimination (as an indicator of tactile spatial acuity). Here, we investigate PL and DD on the torso in the same subjects, using 3×3 arrays of tactile stimulators.

The majority of torso-worn tactile displays developed in the last two decades have commonly adopted miniature affordable vibrotactile stimulators in commercial and experimental frameworks (Arafsha et al. 2015; Karafotias et al. 2017; Garcia-Valle et al. 2018). For instance, Van Erp and colleagues have employed torso-worn vibrotactile displays for use as a pedestrian navigation system (van Erp et al. 2003; Van Erp et al. 2005; Van Erp 2005b; van Erp 2007). Lemmens et al. (2009) developed a wearable vibrotactile jacket to investigate the potential intensification of emotional immersion while participants watched a movie. Garcia-Valle et al. (2018) showed that using a haptic vest, which presented vibration patterns, improves immersion in multimodal virtual reality environments. Other studies have recently employed force stimulators to present collision-type touch stimuli (e.g., force, pressure, compression, etc.) on the participants’ torso (Delazio et al. 2018; Al-Sada et al. 2019; Fadaei et al. 2021). For instance, a force jacket was made of pneumatically actuated airbags to provide strong and variable forces to the torso along with vibrotactile sensations (Delazio et al. 2018). In this line of research, it has been shown that the level of immersion in a virtual environment could be considerably enhanced by presenting ecologically-valid touch feedback (Yoshikawa and Nagura 1997; Lopes et al. 2015; Cao et al. 2018). In addition, the processing of tactile spatial directional cues and notification has been described as more intuitive to participants when using force stimulation rather than vibrotactile as collision touch sensations are a more common haptic experience in daily life. Until now, however, research about tactile spatial discrimination on the torso has focused on performance for manually applied stimuli (Weber 1834; Weinstein 1968; Green 1982; Gibson and Craig 2005) and vibrotactile stimuli (Eskildsen et al. 1969; Jones et al. 2009; Hoffmann et al. 2018). To the best of our knowledge, there is no study investigating tactile spatial perception on the torso for directed force stimuli. Thus, it is not known whether force stimuli are characterized by improved tactile performance (compared to vibrotactile stimuli) in spatial discrimination tasks on the torso. Indeed, force and vibrotactile stimuli present some distinct features; for instance, vibrotactile stimuli are known to spread beyond the limits of the contact area (Cholewiak and Collins 2003), while force stimuli are more focal, and this might lead to better accuracy in spatial discrimination. Here, we measured performance in PL and DD tasks and investigated the spatial accuracy of focal force and vibrotactile stimuli on the torso.

Moreover, previous studies have demonstrated directional anisotropies in tactile localization and tactile spatial acuity for both static pressure and vibrotactile stimuli. These studies reported higher tactile spatial performance along the transverse (limb) axis compared to the vertical axis. In particular, for the PL task on the back, Jones and Ray (2008), using a 4 x 4 array of vibrators, observed that participants were better in the horizontal (87% correct) than vertical direction (68% correct) when using vibratory stimuli. Also, for the DD task, recently, Hoffmann et al. (2018) found that vibrotactile DD accuracy is substantially higher in the horizontal axis compared to the vertical on the lower thoracic region, consistently across three different types of vibrators. Therefore, we also tested for any direction anisotropies in the PL and DD, using force and vibrational stimuli.

In summary, in the present study, we employed two body-conforming torso-based tactile displays (arrays of 3×3 vibrotactile stimulators: Vibrotactile vest; force stimulators: Force vest) and assessed tactile PL and DD on the skin surface of the human upper thoracic area. Using a with-in participant design, we 1) evaluated tactile spatial discrimination (PL and DD) of the upper torso region, 2) examined the association between the results of PL and DD tasks, and 3) compared performance when using vibrotactile and force stimulations. Finally, 4) we searched for directional anisotropy in both tasks and both types of stimulation.

## Materials and Methods

### Participants

A total of 34 healthy participants (17 females, aged between 20 and 36 years, M = 26, SD = 4.2) were recruited for the experiment. All participants were right-handed (assessed via a 12-item Edinburgh Handedness Inventory (Oldfield and others 1971)). Pathological conditions affecting tactile sensitivity (e.g., skin alteration, chronic pain, fractures) were excluded. They provided informed consent and ethical approval that was granted by the cantonal ethics committee in Geneva. All participants received a compensation of 20 CHF/hour for their commitment to the experiment.

### Apparatus

#### Vibrotactile vest

The Vibrotactile vest consists of 9 (3 x 3) coin-shaped, Eccentric Rotating Mass (ERM) vibrators (310-003, Precision MicroDrive; body diameter: 10mm; body length: 3.4 mm; weight: 1.1 gr) with an inter-tactor distance of 60 mm (Fig 1a and Fig 1c). The ERMs are controlled by haptic motor drivers (DRV2605, Texas Instruments) on 5 V (DC), resulting in a vibration frequency of 175 Hz and acceleration of 1.3 G. The haptic motor drivers were controlled with a microcontroller (STM32F407, STMicroelectronics; sampling time of 1 ms) which connects via Bluetooth to a host PC. A custom-made GUI was developed using the Qt platform (free and open-source platform to create GUI) to control vibrators and the experiment flows as well as record participants’ responses (i.e., entered via numeric keypad) along with the experiment. The ERM vibrators were attached to a 20mm-thick foam (Softpur polyurethane foam) using glued-on snap fasteners. Vibrator foam was fixed to a fully-elastic, posture-corrector brace using Velcro straps, allowing the experimenter to change or replace the vibrator foam easily. The Vibrotactile vest covers the whole back, and it is unisex. The front part of the brace includes elastic straps that wrap around the shoulder, chest, and lower back to ensure a snug and secure fit (see Fig 1b). Moreover, the specific load frequency for the ERM vibrators was tested by activating each vibrator while the Vibrotactile vest was firmly fitted to a participant’s torso. The frequency of each vibrator was analyzed using real-time fast Fourier transform analysis (Audio Spectrum Analyzer dB RTA). Our test results showed that the load frequency ranged between 150 and 220, with an average of 175 Hz.

**Fig. 1.**
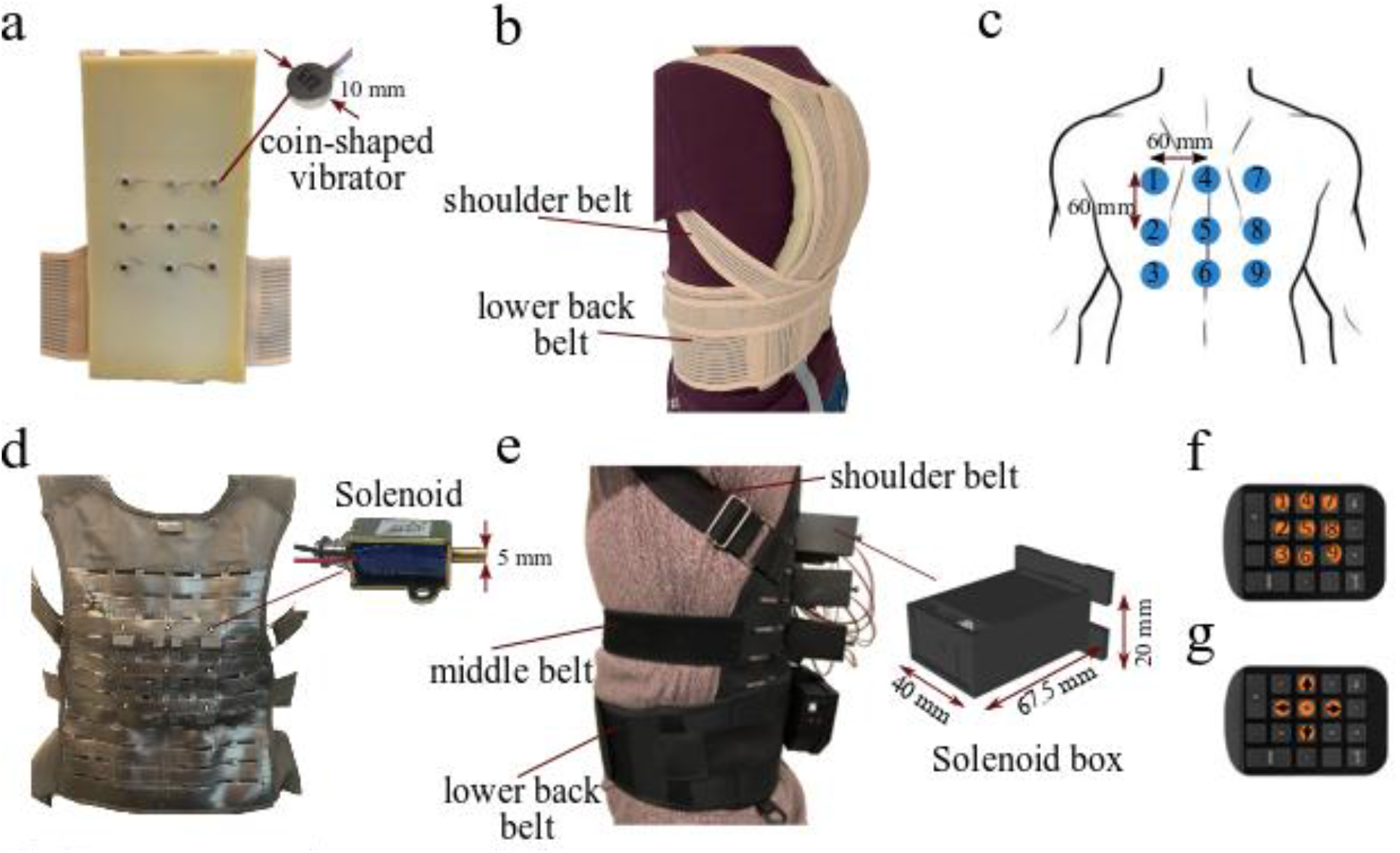
Experimental setup. (a) Interior view of the Vibrotactile vest with 3×3 of coin-shaped ERM vibrators. (b) The Vibrotactile vest on the participant. The vest was firmly fitted on a participants’ body with the lower back and shoulder belts. (c) Arrangement and numbering of stimulations for both Vibrotactile vest and Force vest. (d) Interior view of the Force vest with 3×3 push-pull solenoid actuators. (e) The Force vest on a participant. Three stretchable belts, including shoulder, chest, and lower back belts, firmly fixed the vest on the participants’ torso. Solenoids were placed in a custom-made 3D-printed box. (f) A numeric keypad with marked buttons was used to respond to the PL task. (g) A numeric keypad with marked buttons is used to respond to the DD task.

### Force vest

The Force vest is torso-worn and can apply focal force stimuli to the back. It was designed and prototyped in our previous study (previously named Cogno-vest) (Fadaei et al. 2021). It consists of nine (3 x 3) force stimulators (bi-directional, push-pull solenoid actuators; starting force: 5 N at 12 VDC, shaft length: 5.5 mm; shaft diameter: 5 mm; weight: 39 g), situated with an inter-stimulator distance of 60 mm on the back part of a tailor-made, Y-harness brace (Fig. 1d). To overcome gender-specific morphology, the front part of the brace consists of stretchable straps that wrap around the shoulder, chest, and lower back. The back part is made of polyester nylon with integrated laser-cut loops to support the hardware (Fig. 1e). Each force stimulator is embedded in a customized 3D-printed box, mounted on the back of the brace (see solenoid box in Fig. 1e). The Force vest is thus unisex and can keep stimulators flush against the skin. Arduino Mega 2560 controls the driving of solenoids via Bluetooth to a host PC with a sampling time of 1 ms (similar to the Vibrotactile vest). A custom-made GUI was implemented in the Qt platform to provide a convenient interface to control stimulators with the Force vest and to facilitate the running of the experiment. It also recorded participants’ responses (i.e., entered via numeric keypad). In our earlier study (Fadaei et al. 2021), the force stimulator (solenoid) performance was evaluated under various environmental and parametric conditions (e.g., the effects of the 3D printed box, elastic tips, and stroke length). The results revealed that the realistic amount of force (average) provided by the Force vest is between 0.5 and 0.8 N depending on the stroke length (the force going up as the stroke length increases until 4 mm). The load frequency of the force stimulator was assessed and analyzed in the same way as described for the vibrotactile stimulator. The results revealed that the load frequency ranged between 500 and 1000 Hz, with an average of 650 Hz.

### Experimental design

Participants were exposed to a repeated measure design; participants wore both haptic vests (i.e., Vibrotactile vest and Force vest) and completed both the PL and DD tasks. In the experimental design for the PL task, two factors were manipulated: stimulation type (vibrotactile vs. force) and stimulator location (9 locations on the upper thoracic region). For the DD task, stimulation type (vibrotactile vs. force) and tactile orientation (three different orientation presentations, including horizontal (H), vertical (V), and double activation (DA)) were manipulated as independent within-participant variables.

### Procedure

Participants were randomly assigned in a balanced way to the first experimental session with one of the two vests (Vibrotactile vest or Force vest). Subsequently, they performed both the PL and DD tasks in random order with a break (7mins). After, they changed the vest and were exposed to a second experimental session where they performed the same tasks in a counter-balanced order with respect to the previous session (see Fig. 2).

**Fig. 2.**
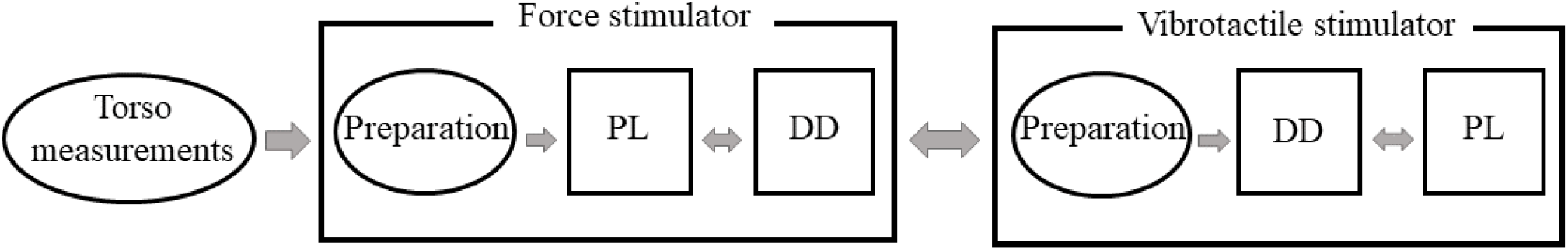
Experiment flow (simplified representation).

At the beginning of the study, participants were asked to wear a thin, fitted white T-shirt. This was done to eliminate any cloth-specific effect. Next, the experimenter performed the torso measurements, which included three measurements, namely torso length (vertical distance between the 7th cervical (C7 vertebra) and the top of the hip bone (iliac crest)), waist circumference (between the belly button and rib cage), and chest circumference (at the fullest part of the bust). Throughout the experiment, participants wore headphones playing white noise to conceal the activation noise generated by the haptic stimulators. The white noise intensity was customized for each participant to have full acoustic isolation.

Prior to each session, the experimenter helped participants to wear the haptic vests correctly. Stimulator arrays were placed centrally on the thoracic regions on the back, starting from the shoulder blades (scapula bones). Each session began with a calibration phase, where the experimenter activated each stimulator (i.e., vibrotactile or force) individually (with a duration of 250 ms and in a random sequence) to ensure that the participant could feel all stimuli by obtaining verbal confirmation. Due to the different nature of stimulators used in the Vibrotactile vest and Force vest, the calibration procedure was different for the two vests. For the Vibrotactile vest, in case of failure in perceiving the vibrotactile stimuli, the experimenter improved the perception by better fitting the vest on the participant’s torso. On the other hand, for the Force vest, the experimenter manually adjusted the force stimulator (solenoid) position at different stroke lengths until receiving verbal confirmation from the participant that they felt mechanical touch by each stimulator (more details can be found in (Fadaei et al. 2021)). The calibration tasks for the Vibrotactile and Force vest lasted approximately 5 and 10 minutes, respectively. Participants completed a training session, similar to the main task but shorter (around 1 minute). During the training session, participants learned how to respond to tactile stimulation on their back with the corresponding keypad (see Fig. 2).

In the PL task, a series of discrete tactile stimuli were applied to the participants’ back. They were instructed to indicate the location of the perceived stimulus, i.e., the position out of the total of 9 locations where the tactile cue had been applied (9 alternatives forced choice, 9-AFC). Participants responded by specifying a number that corresponded to the stimulated location by pressing a numeric keypad, as indicated in Fig. 1f). Participants did not receive performance feedback during the task. To reduce task complexity, they were asked to keep the keypad so they could see the buttons in the same order as the stimulators numbering on the back. In each trial, stimulators were activated for 250 ms with a random inter-trial interval of 2000±250 ms. Tactile point activations (each of 9 different locations) were repeated 20 times, resulting in a total of 180 trials. To reduce potential fatigue, the task was divided into two blocks, each one including 100 trials and lasting 3 minutes. There was also a short break of around 2 minutes between two blocks.

In the DD task, participants received two consecutive stimuli, and they were asked to determine whether the second was to the right, left, above, or below the first one or whether the same location was stimulated twice (5-AFC). Considering the arrangements of stimulators on the vests (3 x 3), shown in Fig. 1c, there are a possible 12 different horizontals (H; i.e., along with transverse axis) and vertical (V; i.e., along with longitudinal axis) presentations, and nine DA of the same stimulator. Each of the orientational combinations was repeated five times, and the DA condition was repeated six times (to assess the same number of repetitions per condition), resulting in a total of 174 trials. The location of the first stimulus and the relative position of the second were randomly arranged. Participants responded via a standard numeric keypad with five marked buttons (see Fig. 1g) corresponding to the five possible response options, and the software recorded their responses. Similar to the PL task, stimulators were turned on for 250 ms with an inter-stimulus interval of 50 ms. The inter-trial interval was altered randomly in the range of 2000±250 ms. To avoid fatigue, the task was divided into two blocks of 87 trials, and each block lasted 4 minutes. There was a short break of approximately 2 minutes between two blocks.

### Statistical Analysis

All analyses were performed in R (R Core Team 2020) running in the RStudio environment (RStudio Team 2020). In the DD task, two participants had very low accuracy across the vests. Those data were excluded from DD analysis as, presumably, the two participants did not understand the DD procedure correctly. Thus, those analyses that involved DD data only included data from 32 participants.

For the between-stimulus comparison, the average accuracy (in percentage) of each task was considered as the response. A two-tailed paired sample t-tests were used to assess whether accuracy differed significantly between two stimulations. The chance level for PL and DD tasks were estimated at 11.11% and 20% since there were 9 and 5 possible response types in each trial, respectively. One-sample t-tests were used to compare the accuracy with the chance level.

To better understand the PL accuracy and its variation on the upper thoracic regions of the back, further analysis was conducted by considering accuracy (in percentage) at each location (location 1 to 9) as the response. To investigate the participants’ ability to identify the stimulator’s location along with the vertical (column) and horizontal (row) axis on the back, data were collapsed across columns (upper, middle, and lower columns) and rows (right, middle, and left rows) stimulators. Thus, a linear mixed-effect model was performed to assess the effect of vibrotactile vs. force by considering stimulation rows, columns, and interactions between them as fixed effects and the participant as a random effect, accounting for between-subject variability.

To explore orientational biases in the PL task, the number of localization errors were computed at each location for both vests. Mislocalization data were collapsed into adjacent (N_Adjacent_: confusion with stimulators that are situated one gap away from the target) versus nonadjacent biases (N_Nonadjacent_: confusion with stimulators that are situated at more than one gap away from the target), and horizontal (N_H_: number of errors made along with horizontal axis) versus vertical (N_V_: number of errors made along with longitudinal axis) biases. As inferred with the Shapiro-Wilk test of normality, mislocalization data significantly deviated from the normal distribution. Therefore, the Wilcoxon signed-rank test was employed to investigate the effect of adjacent and orientational biases.

To investigate the orientation-dependent effect in DD results, mean accuracies for horizontal (DD_H_) and vertical (DD_V_) trials were considered as the response. Then, a linear mixed-effect model was used by considering stimulation (vibrotactile vs. force), tactile orientation (H vs. V), and their interaction as fixed effects and subject as a random effect.

All post-hoc comparisons were conducted using the Tukey HSD test. Pearson correlation coefficient was calculated to investigate the relationship between variables, and the p-values were corrected for multiple comparisons (Bonferroni correction). In all analyses, significance was reported for p-values smaller than 0.05.

## Results

### Overall accuracy

Overall accuracy results are represented in the Fig. 3a. In both tasks, participants were significantly better than chance level (PL: vibrotactile: t(31) = 34.78, p < 0.001; force: t(31) = 29.9, p < 0.001; DD: vibrotactile: t(31) = 25.77, p < 0.001; force: t(31) = 17.77, p < 0.001). Performance in the PL task was significantly higher with the vibrotactile stimulation (M = 60.70%, SEM = 2.32) versus the force stimulation (M = 54.6%, SEM = 1.95; t(33) = −2.92, p = 0.006). For the DD task, no significant accuracy difference was found in between the two stimulations (vibrotactile: M = 71%, SEM = 1.98%; force: M = 67.7%, SEM= 2.69%; t(31) = −1.54, p = 0.13).

**Fig. 3.**
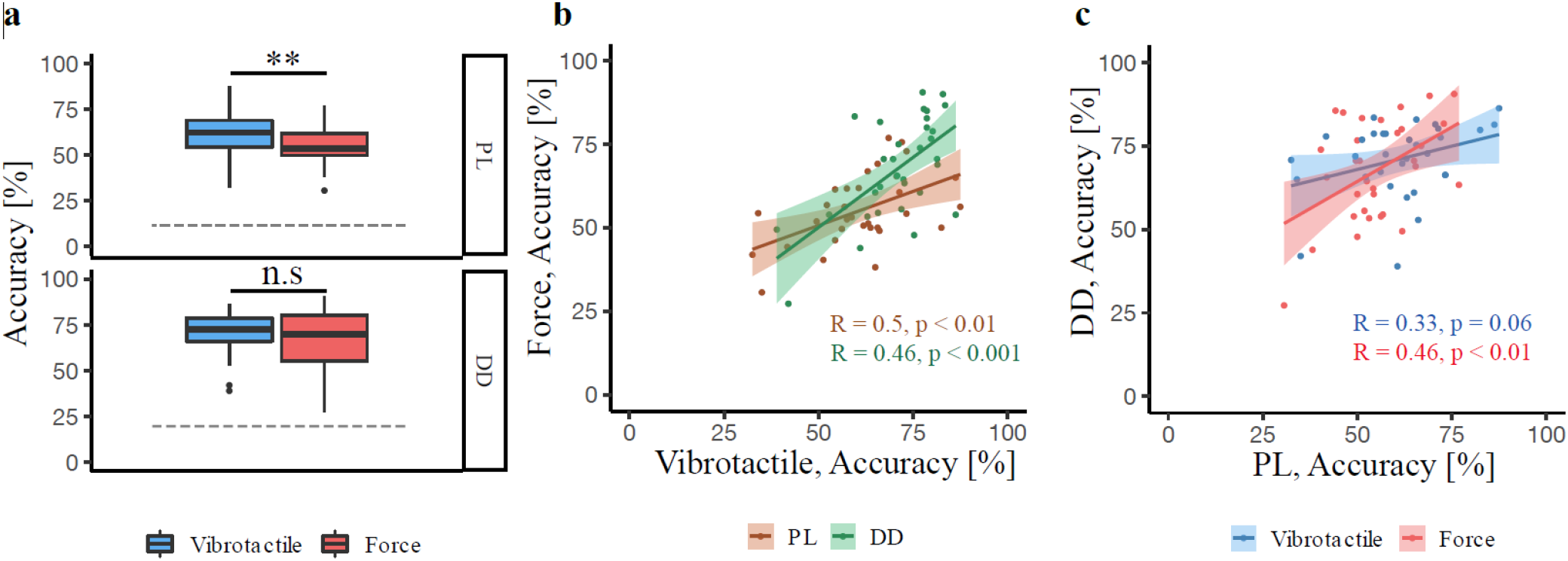
Overall accuracy results. (a) Box plot of overall accuracy for two stimuli across tasks. PL accuracy was significantly higher with vibrotactile stimulation, while no difference was found between DD accuracies of two stimulations. Gray dash-lines represents the chance level. Each box plot shows the median (50th percentile; dark bar), values to the 1.5 interquartile range (whiskers), 25th to 75th percentile range (box), and outliers (*p < 0.05 and **p < 0.01). (b) Scattered dot plot and Pearson correlation between accuracies with vibrotactile and force stimulations across tasks. There are positive correlations between the accuracies of two simulators for both tasks. Each point represents data from a single participant, and shaded areas show the 95% confidence interval for the regression line. (c) Scattered dot plot and Pearson correlation analysis between PL and DD accuracies of two stimulations. There is a significant positive correlation between PL and DD accuracies for force stimuli (in red) and not vibrotactile stimuli (in blue).

The tactile performance was found to correlate between tasks (PL, DD) and stimulations (vibration, force). Thus, participants’ performance with vibrotactile stimulation significantly correlated with participants’ performance using force stimulation in the PL task (R = 0.50, p = 0.002; brown line in Fig. 3b) and in the DD task (R = 0.61, p < 0.001; green line in Fig. 3b). This was also found when comparing the two tasks, revealing significant correlations between the two tasks for force stimulation (R = 0.46, p = 0.007; red line in Fig. 3c), but for the vibrotactile stimulation there was only a trend towards a significant correlation (R = 0.33, p = 0.060; blue line in Fig. 3c).

### PL task

The range of localization accuracies across both types of stimulation was 52.0-71.7% for vibrotactile and 37.5%-65.1% for force stimulation, respectively. The mixed-model showed a significant main effect of stimulation (F(1, 573) = 11.22, p < 0.001), stimulation row (F(2, 573) = 28.52, p < 0.001), and stimulation column (F(2, 573) = 8.22, p < 0.001); none of the interaction terms were significant. In line with the overall accuracy findings, participants’ PL accuracy was significantly higher for the vibrotactile versus force stimulation (t(561) = −3.34, p <0.001). Fig. 4a shows that, whereas PL performance in peripherial areas did not significantly differ (right-left columns: t(573) = 0.90, p = 0.6), performance was significantly more accurate for stimulations in peripheral than midline columns (right versus middle column: t (561) = 2.97, p = 0.009; left versus middle columns: t (561) = 3.87, p < 0.001, Fig. 4a). In addition, as it is shown in the Fig. 4c, stimulations located in the middle row were more accurately perceived compared to the upper (t(573)= −6.97, p < 0.001) and lower rows (t (66) = 5.79, p <0.001). No significant difference was found between accuracies of the upper and lower rows (t(573) = 1.18, p =0.4).

**Fig. 4.**
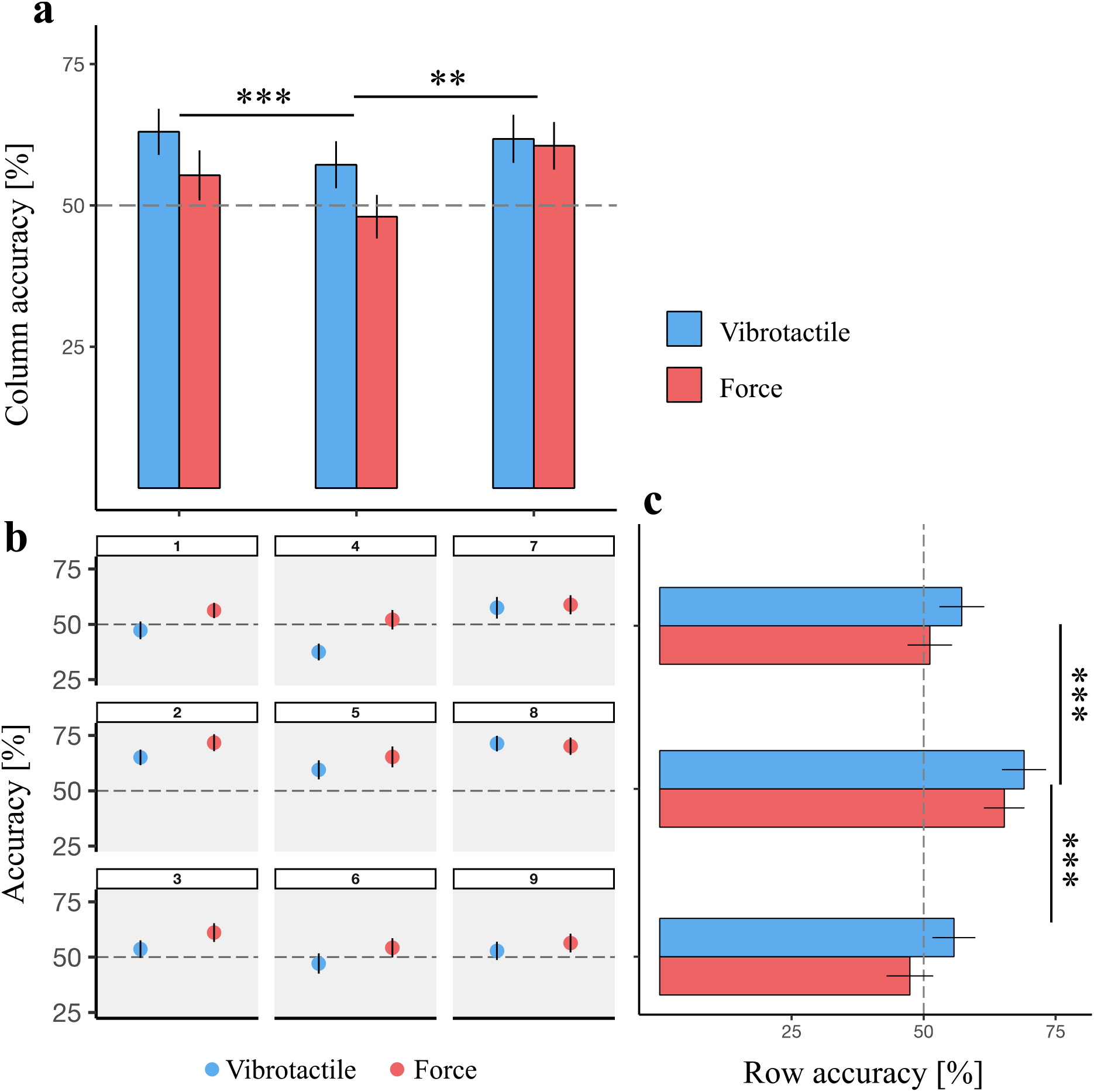
(a) Mean PL accuracies at three columns in the array for both stimulations. (b) Mean PL accuracies at nine stimulation landmarks in a 3 x 3 array for both stimulation types. (c) Mean PL accuracies at three rows in the array for both stimulations. The dashed line shows the 50% threshold, and error bars illustrate the standard error of the mean (SEM) (* p < 0.05, ** p < 0.01, *** p <0 .001).

Table 1 lists localization errors in the PL task. For both stimulations, the majority of such errors were characterized by a mislocalization to an adjacent location (adjacent versus nonadjecent location: vibrotactile: z = 595, p < 0.001; force: z = 595, p < 0.001). Analyzing whether there was an axis along which localization errors predominated, we found that the number of horizontal errors was significantly lower than vertical errors, across stimulations (vibrotactile: z = 5, p < 0.001; force: z = 5.1, p < 0.001).

**Table 1.**
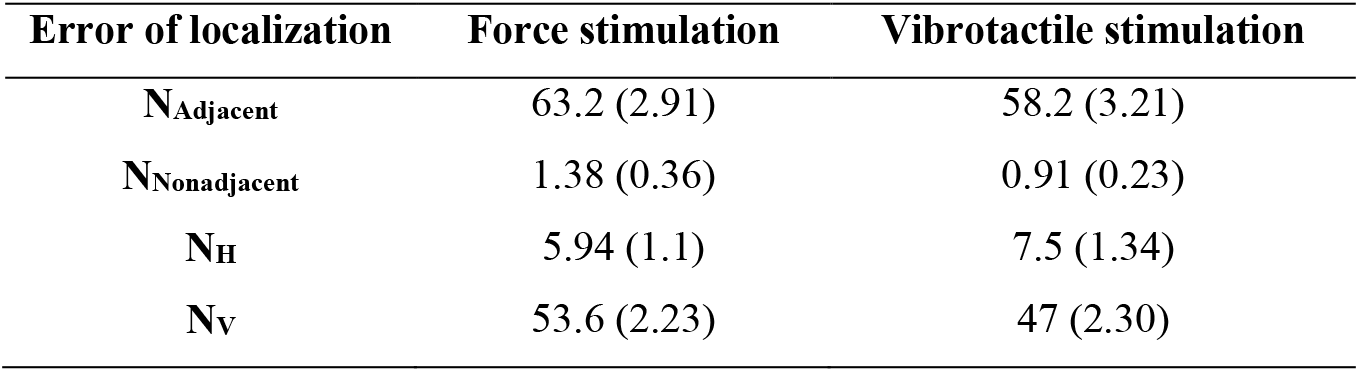
Means of different tactile localization errors (standard error of the mean) for two vests.

| Error of localization | Force stimulation | Vibrotactile stimulation |
| --- | --- | --- |
| N<sub>Adjacent</sub> | 63.2 (2.91) | 58.2 (3.21) |
| N<sub>Nonadjacent</sub> | 1.38 (0.36) | 0.91 (0.23) |
| N<sub>H</sub> | 5.94 (1.1) | 7.5 (1.34) |
| N<sub>V</sub> | 53.6 (2.23) | 47 (2.30) |

We further compared the number of localization errors between the two types of stimulation for different categories of error (i.e., adjacent vs. nonadjacent, and H Vs. V). Results showed no significant difference in the number of localization errors between the two stimulations (all p>0.05).

### DD task

Investigating the effect of tactile orientation and of stimulation type on the accuracy in the DD, we found a significant main effect for tactile orientation (F(1, 93) = 37.33, p < 0.001) and a significant interaction between the stimulation and tactile orientation (F(1,93) = 8.68, p < 0.001). Post-hoc analysis showed that participants were more accurate for trials presented along the horizontal axis only when using vibrotactile stimulation (t(93)= 6.76, p < 0.001; Fig. 5) (this effect was absent for force stimulation (t(93) = 2.24, p = 0.12)).

**Fig. 5.**
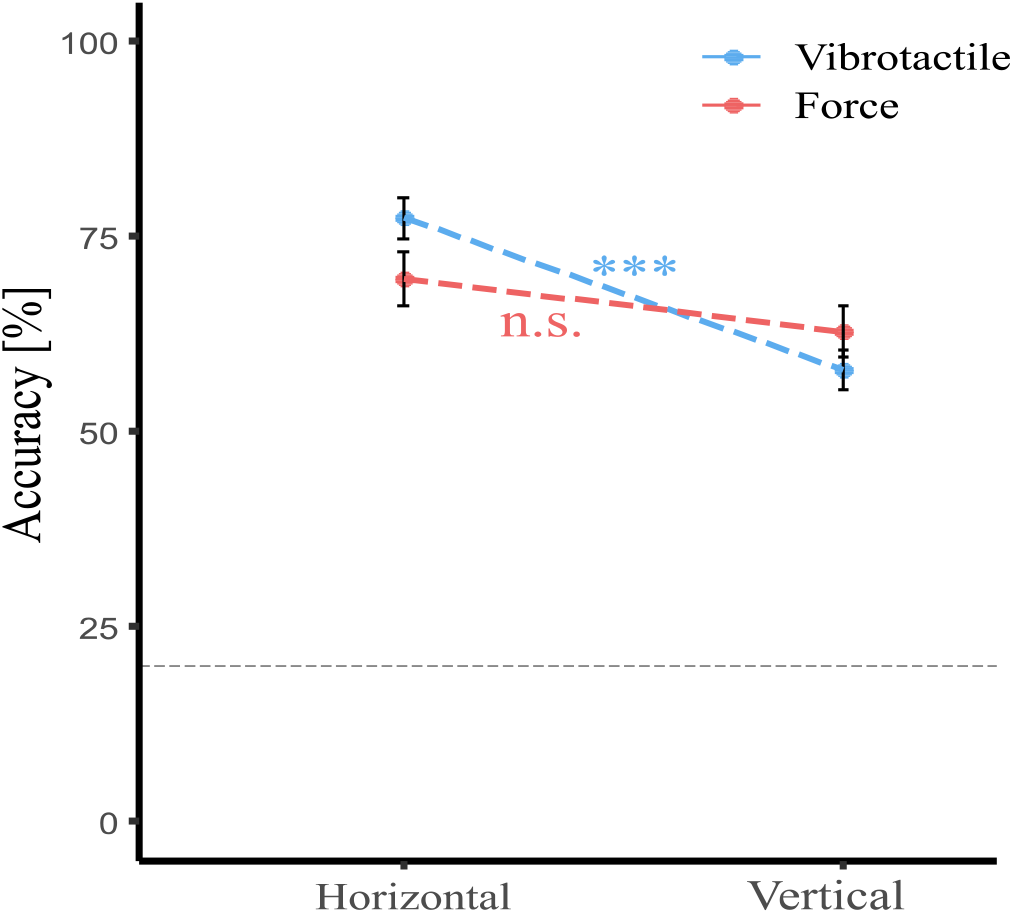
DD accuracies with vibrotactile and force stimulations. The dash-line shows the chance level of 20%. The error bars show the standard error of the mean (SEM) (*** p <0 .001; n.s.: No significant).

We assessed correlation coefficients separately for vertical and horizontal levels to explore any potential relationships between orientation-related effects in DD accuracy (worse along the vertical axis) and localization errors in PL (higher vertical errors). The only significant correlation was found for the vibrotactile stimulation in the vertical axis (DD_V_-N_V_: R= −0.36, p = 0.040; Bonferroni corrected p-value), revealing that those participants who had higher vertical errors (N_V_) in the PL task they also had low DD performance in discriminating vibrotactile stimulation in vertical axis.

## Discussion

In the present study, we adapted two automatized tactile perception paradigms, namely tactile PL and tactile DD task, and measured tactile spatial discrimination on the human back. More specifically, we evaluated tactile perception over the thoracic region, using two custom-made haptic interfaces consisting of either vibrotactile or force stimulators. We found that PL and DD accuracy were slightly higher with vibrotactile stimulations than those with force stimulations. Using a within-participant design, we further demonstrate that tactile performance generalizes across tasks (PL and DD) for force stimulations but not for vibrotactile ones. Furthermore, we observed directional anisotropies in both tasks characterized by better performance for horizontal directions.

### Overall accuracy

We observed an overall PL accuracy of 60.7% for vibrotactile and 54.6% for force stimulation in the thoracic region on the back. In the DD task, we reported an overall accuracy of 71% for vibrotactile and 67% for force stimulation. Our PL results (accuracy) are in line with earlier studies that employed a two-dimensional array of vibrators reporting PL accuracy around 60% when stimuli were applied to the participants’ lower back (Cholewiak and McGrath 2005; Jones and Ray 2008). Considering the DD task, our vibrotactile accuracy was lower than in a recent study (Jóhannesson et al. 2017), where the accuracy of 91% was found, even if the inter-stimulator distance of 30 mm was smaller than in our study (60 mm). However, the latter authors used a 3-AFC DD task (via 3×3 vibrotactile array), which has a lower degree of difficulty compared to the 5-AFC used in the current study.

### Vibrotactile vs. force stimulation: correlation and comparison

We observed a moderate, positive correlation between performance for both stimulators across tasks (see Fig. 3c), suggesting that the spatial discrimination processes for both stimulation modalities, as tested in the present study, rely on similar perceptual mechanisms. Load frequencies of the vibrator (175 Hz) and force stimulator (600 Hz) fall in the frequency range of the Pacinian corpuscle sensitivity (100-1000 Hz) (Vallbo et al. 1984). The relatively low tactile spatial discrimination rate (around 60%) reported for both stimulations and across tasks further points to this type of subcutaneous mechanoreceptor, with its comparatively large receptive fields. In comparison, non-Pacinian receptors embedded in the glabrous skin of the palm or fingers have shown higher tactile spatial sensitivities (Weinstein 1968).

While accuracy was higher for the vibrotactile than for the force stimulation across both tasks (mean difference, PL: 6.1%; DD: 3.3%), this difference was only significant for the PL task. We fixed the spatial parameters (i.e., body site, inter-stimulator distance) and temporal parameters (i.e., burst duration and inter-stimulus interval) between the two stimulations as they may have a profound effect on both PL and DD results (Cholewiak et al. 2004; Van Erp 2005a; Jóhannesson et al. 2017). Nonetheless, we observed greater accuracy when using vibrotactile stimulation, suggesting that other stimulus-properties account for this discrepancy. These could include physical features (frequency, intensity, mass), contact area, the direction of movement with respect to the skin, and the amount of surface wave created by activating the motor. We here observed a higher accuracy with vibrotactile stimulations, which had a lighter weight (1.1 gr vs. 39 gr), slightly lower acceleration (1.3 G vs. 1.65G), and lower load frequency (175 Hz vs. 650 Hz). Considering that the combination of mass, acceleration, and frequency of the tactile stimulation contributes to the perceived force, our results corroborate previous studies (Gibson and Craig 2006; Hoffmann et al. 2018), suggesting that the effect of physical parameters is not sufficient to account for the performance in the PL and DD tasks. For the effect of frequency, it has been found that increases in frequency above 80 Hz (i.e., Pacinian corpuscle) did not result in higher accuracy (Cholewiak et al. 2004; Cholewiak and McGrath 2005; Hoffmann et al. 2018). Another possible explanation for the higher PL and DD accuracy with vibrotactile stimulations might also be that the contact area of the vibrotactile stimulator was twice as large as the one chosen for the force stimulator (vibrotactile: 314 mm^2^, force: 78.5 mm^2^). While the effect of the contact area on the PL/DD has not been systematically investigated, it would seem that due to the spatial summation of afferent signals from Pacinian corpuscles, vibrotactile thresholds (above 50 Hz) decrease as the stimulator area increases (Verrillo 1963; Gescheider et al. 2010), presumably enhancing the perceptual capabilities. In addition, we here observed that PL and DD accuracy were higher with the vibrotactile stimulator (coin-shaped vibrators), which generates motions parallel to the skin’s plane, compared to force stimulators that generate force perpendicular to the skin. This observation may relate to previous findings of Hoffmann et al. (2018), who found that DD accuracy was higher with vibrators that generate motion parallel to the skin (as in our study) compared to the perpendicular to the skin as it provides stronger surface waves traveling on the skin (see Hoffmann et al. 2018 for more details). Future work is needed to investigate how tactile perception on the back depends on these different mechanisms.

### Association between PL and DD results

In the present study, we found a positive correlation between the PL and DD performances for the force stimulations but not for the vibrotactile stimulations. Although prior studies often directly compared the results of tactile localization (PL) with tactile spatial acuity (e.g., DD), our results suggest that extending the results of PL to DD (or the other way around) depends on the type of tactile stimuli and may not hold for vibrotactile stimulation (at least in the present study). The absent correlation between the PL and DD tasks with vibrotactile stimulations suggests that distinct and, presumably, stimulation-related driving factors are involved in the discrimination process of the vibrotactile DD task as tested by us. As discussed below, the DD task with the vibrotactile stimulation may more heavily depend on and vary with the viscoelastic properties of the participants’ skin.

### PL Task

We observed that the PL accuracies for both stimulations vary over the skin surface of the back in the thoracic region (vibrotactile: 52%-71.68%, force: 37.47%-65.08%). This observation may partly be explained by garment conformity but may also reflect differences in mechanoreceptor density on the torso. The present PL performance is in line with previous studies that reported variation in tactile PL (for both static pressure and vibrotactile stimuli) across the skin surface for different body parts, also including the back (e.g., forearm, abdomen, lower back, palm, thigh, etc.) (Cholewiak and Collins 2003; Oakley et al. 2005).

### Lower PL accuracy on the spine compared to the peripheral area

Concerning the PL performance on the spine, previous studies have yielded mixed results and further depended on the physical arrangement of stimulators. Some studies that used a one-dimensional array of vibrators supported the enhancement of PL in midline regions (close to the spine) as an anatomical body reference (i.e., as being related to the joints of the body) (Boring 1942; Cholewiak and Collins 2003; Cholewiak et al. 2004; Van Erp 2005a; Jones and Ray 2008). Others, however, did not report improved PL in midline regions when using two-dimensional arrays (Lindeman and Yanagida 2003; Jones and Ray 2008). In the present study, we observed that the PL accuracy was lower for stimulators in the midline area (column 2) than those located peripherally (columns 1 and 3). These results match the findings of previous multi-dimensional setups, suggesting that by adding dimension to the tactile display (e.g., two-dimensional array), no midline advantage is observed. Compatible with this account, earlier studies on the PL for the forearm also found increased accuracy at the edges of the arm compared to those in the center (Oakley et al. 2005; Chen et al. 2008). Based on the present data of enhanced performance for lateral stimulations, we speculate that perceptual and attentional mechanisms related to lateralized stimulations may boost performance. However, more work is needed to specifically test this hypothesis and its comparison with enhanced midline performance. We also note that structural aspects such as the higher curvature in the spinal area compared to lateralized torso locations and the consequent poorer fit of the vest and stimulators around the midline may also play an important role.

### Directional anisotropy

In the present study, we observed directional anisotropy in PL performance in both tasks, as participants made considerably fewer localization errors in the horizontal than vertical axes (see Table 1). Using a 4×4 array of vibrotactile on the back, Jones and Ray (2008) also showed that participants were less accurate in identifying the correct row of activation than the column. These observations are also in line with earlier PL studies that reported systematic biases in the vertical axis on the skin surface of the palm, thigh (Sofia and Jones 2013), and arm (Oakley et al. 2005; Chen et al. 2008; Sofia and Jones 2013). In addition, we, here, found that the level of anisotropy in PL performance was comparable between the two tasks, suggesting that comparable mechanisms are involved in the spatial discrimination process for both stimulations. One possible explanation, as discussed above, is that the torso’s lateral sides function as a perceptual reference point (as located to the endpoints of the stimulus range) affording higher accuracy in the horizontal direction. It has also been proposed that receptive fields of afferent fibers and/or neurons in the spinal cord and somatosensory cortex are oval-shaped and elongated along the longitudinal-vertical axis (Cody et al. 2008). Thus, one may speculate that stimulators in the horizontal axis would more likely activate separate adjacent mechanoreceptors, leading to fewer localization errors along with the horizontal axis. However, these argumentations do not seem to apply for our observation concerning DD results as anisotropy was only seen in the results with vibrotactile stimulations: higher DD accuracy for vibrotactile stimulation in the horizontal axis than vertical. Our observation for vibrotactile stimuli is consistent with the findings of Hoffmann et al. (2018), who recently found the superior DD accuracy for vibrotactile stimulations presented horizontally on the lower thoracic region, across different types of vibrators. We argue that anisotropy in DD results is mainly influenced by the amount of surface waves created by force and vibrotactile stimulations and how they spread on the skin. In contrast to focal force stimulation, vibrotactile stimuli spread beyond the contact area in the form of surface waves. As the skin is a highly viscoelastic tissue, its mechanical properties highly impact the spread of the surface wave from the vibration source. Some direct human and animal evidence suggested that skin stiffness is anisotropic with higher stiffness along with the vertical axis than horizontal. Therefore, surface wave from a vibrating source may propagate further along with the vertical axis and hence excite mechanoreceptors some distance from the cite of the stimulation, which makes difficult the recognition of the stimulation direction. (e.g., in our case, up or down).

### Wearability challenge of torso interfaces

While waist-worn tactile displays have been widely used in many recent studies (Jones and Ray 2008; McDaniel et al. 2008) to investigate vibrotactile spatial discrimination (PL and DD) at the lower torso, very little research has assessed the tactile spatial resolution of the upper thoracic torso. One complication has been the large morphological differences in the human torso area (within and between-participants’ variability). Hence, forming and fitting the torso-worn haptic display to the participant’s body is a challenge in assuring consistent stimulus application (Mortimer and Elliott 2017). In our previous study (Jouybari et al. 2019), in spite of using gender-specific chest-belts, containing 3×2 vibrators (horizontal spacing of 15cm and vertical spacing of 8 cm), we obtained poor localization accuracy of 30.7% (on average) stemming in the impaired interface design. We noted that the ideal interface for the upper torso should be sufficiently adjustable to guaranty correct fit, firm support, and free-breathing for users even during movements despite the huge morphological variations in the torso area. Therefore, we, here, developed and employed two unisex, body-conforming, torso-based tactile displays. We further provide quantitative and practical information to designers of torso-based tactile interfaces.

### Study limitations

The present study has several limitations. Although we evaluated and compared tactile spatial discrimination using dynamic force (push-pull solenoid) and vibrotactile (coin-shaped ERM) stimulators, we only controlled for the temporal and spatial parameters between the two stimulators. However, there are other parameters that we did not control with the present systems, such as physical parameters, contact area, or spread of vibration waves. Therefore, the generalization of these results to other setups have to be taken with caution. Future experimental investigations are needed to systematically investigate the effect of such individual actuator properties on spatial discrimination, which was beyond the scope of the current study. Second, the design and control of force simulators are more problematic than vibrotactile stimulators. Thus, force stimulators require actual contact with the human skin to be perceived; moreover, as reported in our previous study with a force vest (Fadaei et al. 2021), the solenoid’s impact force might change in the range of 0.5-0.8 N, depending on the stroke length. Such effects could lessen the quality of force perception resulting in lower tactile perception with force stimulators. Future work may monitor the uniformity of the perceived intensity across the array of actuators by employing an objective calibration procedure, where the participants are asked to rate the intensity of each individual actuators. Finally, for the field to advance, the same experimental procedures and tasks have to be applied across participants, conditions, and different research groups, further empowered by the application of psychophysical methodology.

## Conclusion

Collectively, although previous studies investigated the spatial discrimination of vibration stimuli on the back, our study is the first to investigate both localization and tactile direction discrimination of the upper thoracic spine for two different types of dynamic mechanical stimulations (vibrotactile and force) in a large group of healthy participants. Our findings suggest that designers can use force stimulators to design the torso-worn tactile interface to provide more ecological touch feedback with the (almost) similar level of tactile spatial discrimination accuracy as observed in widespread vibrotactile interfaces. We also noted that overall accuracy with both stimulations is still relatively low (around 60%), indicating that further technological improvements are required to improve torso-based tactile communication systems. Apart from technological advancement, we might speculate that long-lasting training might improve performance. Alternatively, we suggest taking advantage of multisensory-training protocols (i.e., a combination of tactile with auditory, visual, or vestibular) instead of unisensory protocols to produce greater and more efficient learning (Ghazanfar and Schroeder 2006; Shams and Seitz 2008; Proulx et al. 2012). Furthermore, our findings provide new insights into the association between the results of the PL and DD tasks on the torso, indicating that the generalization of the PL results to DD is only valid for focal force stimulations and not for vibrotactile ones which spread further away. These results suggest that studies using vibrotactile interfaces should ideally measure spatial discrimination with different measures in parallel to estimate the actual discrimination accuracy.

## Supporting information

Investigating the effect of gender and torso sizing on tactile spatial discrimination

## Availability of data and material

The datasets generated during and/or analyzed during the current study are available from the corresponding author on reasonable request.

## Declarations

### Compliance with ethical standards

The experimental protocol was approved by the cantonal ethics committee in Geneva and followed the ethical standards laid down in the Declaration of Helsinki.

### Consent to participate

Informed consent was obtained from all individual participants included in the study.

### Open access

