## Supplementary material for "Tactile spatial discrimination on the torso using vibrotactile and force stimulation": Investigating the effect of gender and torso sizing on tactile spatial discrimination

Olaf Blanke
Bertarelli Chair in Cognitive Neuroprosthetics, Center for Neuroprosthetics & Brain Mind Institute, School of Life Sciences, Campus Biotech, Swiss Federal Institute of Technology (EPFL), 1012 Geneva, Switzerland

### Supplementary materials

*Investigating the effect of gender and torso sizing on tactile spatial discrimination*

*Statistical analysis*

We investigated the effects of the fitting and gender variabilities on the DD and PL performance results. Participants' torso sizings (i.e., torso length, waist circumference, and chest circumference) were measured before the experiment (explained in the procedure section). For the PL task, the overall PL accuracy was considered as the response. A linear mixed-effect model was used to assess the significant difference between the two stimulators by considering the stimulator type as a fixed factor, gender as a control variable, torso length, chest circumstances, and waist circumference as covariate and subject as a random effect.

For the DD task, the overall DD accuracy was considered as the response. A linear mixed-effect model was also used to assess the significant difference between the two stimulators by considering the stimulator type as a fixed effect, gender as a control variable, and torso length, chest circumstances, and waist circumference as covariate variables and subject as a random effect.

*Results*

Statistical analysis showed that the PL accuracy was significantly higher with vibrotactile stimulators ($F(1, 33) = 8.55$, $p < 0.01$). Furthermore, we found a significant effect for the chest circumference ($F(1, 29) = 6.98$, $p = 0.01$); suggesting that the PL accuracy improved by increasing the participants' chest circumference (see Fig. 1). This observation most likely further supports our argument that participants used torso edge as reference points with which stimuli can be associated. However, the effect of other covariant factors was insignificant (both $p > 0.2$). We also did not find any significant effect for the gender on the PL results ($F(1, 29) = 0.09$, $p = 0.77$).

For the DD task, statistical analysis revelaed no significant effect of the stimulator type ($F(1,33) = 2.81$, $p =0.1$). There was no significant effect for covarient factors (Torso length: $F(1, 29) = 3.84$, $p = 0.06$; waist circumference: $F(1, 29) = 0.11$, $p = 0.73$; chest : $F(1, 29) = 2.85$, $p =0.1$)  neither for the gender factor ($F(1, 29) = 0.12$, $p =0.73$).

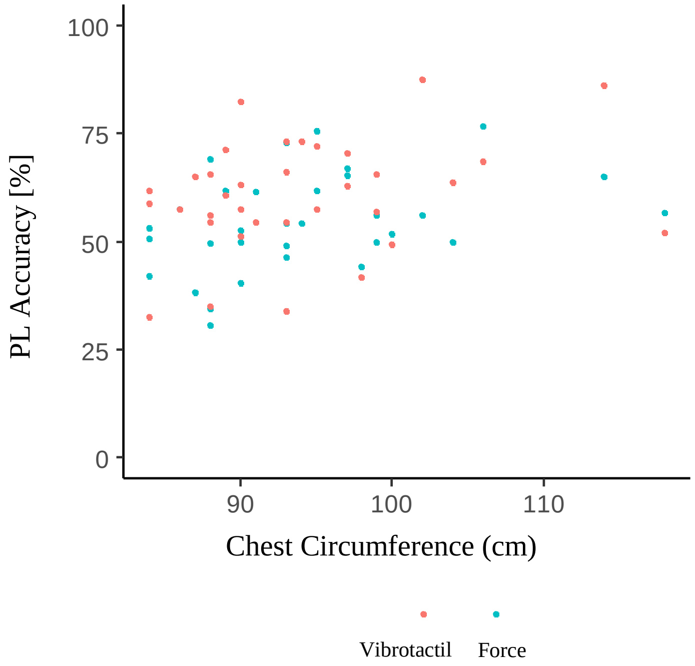


**Fig. 5** PL accuracy as a function of chest circumference (cm). Each point shows the average PL accuracy of one participant with one of the vests.
